# Defocus-dependent Thon-ring fading

**DOI:** 10.1101/2020.10.12.336214

**Authors:** Robert M. Glaeser, Wim J.H. Hagen, Bong-Gyoon Han, Richard Henderson, Greg McMullan, Christopher J. Russo

## Abstract

The brightness of modern Schottky field-emission guns can produce electron beams that have very high spatial coherence, especially for the weak-illumination conditions that are used for single-particle electron cryo-microscopy in structural biology. Even so, many users have observed defocus-dependent Thon-ring fading that has led them to restrict their data collection strategy to imaging with relatively small defocus values. In this paper, we reproduce the observation of defocus-dependent Thon-ring fading and produce a quantitative analysis and clear explanation of its causes. We demonstrate that a major cause is the delocalization of high-resolution Fourier components outside the field of view of the camera. We also show that it is important to make a correction for linear magnification anisotropy, even if it is quite small, before circular averaging of the Thon rings, as is also true before merging data from particles in many orientations. Under the conditions used in this paper, which are typical of those used in single-particle electron cryomicroscopy, fading of the Thon rings due to source coherence is negligible. The principal conclusion is that much higher values of defocus can be used than is currently thought to be possible. This increased understanding should give electron microscopists the confidence to use higher amounts of defocus to allow, for example, better visibility of their particles and Ewald sphere correction.

## INTRODUCTION

Fourier transforms of electron-microscope images of weak-phase objects are modulated by the sine of the wave-aberration function of the objective lens. The contributions to the wave aberration that are normally discussed, as a first approximation, include defocus, astigmatism, and spherical aberration. The resulting modulation of the Fourier transform of an image is referred to as the (phase) contrast-transfer function (CTF), and its manifestation in the power spectra of images of amorphous carbon films produces “Thon rings” (Thon, 1966).

The envelope of the modulation decreases in amplitude at increasing spatial frequency due to multiple factors. The envelope function is normally thought of as being the product of two or more component envelope functions, one due to imperfect spatial coherence and another due to imperfect temporal coherence (Wade and Frank, 1977). Imperfect spatial coherence causes defocus-dependent Thon-ring fading, whereas imperfect temporal coherence causes Thon-ring fading that is independent of defocus. The overall “instrumental” envelope function is often modeled as a Gaussian fall-off.

The defocus-dependent envelope function due to limited spatial coherence has traditionally been the one that is of major concern in high-resolution electron microscopy of weak-phase objects. This historical concern still discourages many electron cryo-microscopy (cryo-EM) users from recording images at high defocus – see Figure 2C in (Cheng, 2015), for example, even though doing so would make it easier to recognize particles and extract them from images. In reality, the spatial coherence that is routinely achieved with a Schottky field emission gun (FEG), especially under the low-dose imaging conditions used for single-particle cryo-EM (Russo and Henderson, 2018a; Börrnert et al., 2018), is high enough that its associated envelope function is no longer a concern, even at resolutions as high as 2 Å.

There are other effects, however, that can produce what might be called “anomalous”, defocus-dependent envelope functions, in the sense that they are not caused by imperfect spatial coherence. In this paper, we find that delocalization of high-resolution Fourier components outside the field of view of the camera (de Jong and Van Dyck, 1993) is currently the main factor that is significant. We here distinguish two categories of delocalization-associated envelope functions.

In the first category, which is the most relevant, the amplitude of CTF oscillations decreases when half of the high-resolution information, at a given spatial frequency, is delocalized – in the form of a single-sideband image – outside of the field of view. Naturally, this occurs first for particles located close to the perimeter of the image. The remaining (Friedel-related) half of the information then contributes an unmodulated, focus-independent contribution to the power (square of the amplitude) of the Fourier transform of the image.

Although Thon-ring modulation is absent in the single-sideband case, the average value of the background-subtracted power spectrum is not expected to go to “zero”, by which we mean a baseline value due to the power spectrum of additive noise. Instead, the remaining, unmodulated power spectrum is expected to fall only to half that of the average power in the modulated, or double-sideband case. This is clearly a behavior that is not explained by imperfect spatial coherence.

In practice, the value of the remaining, unmodulated power spectrum may appear to remain equal to the average, rather than half the average, of the power in the double-sideband case. This is because compensating information, which unfortunately is of no use, may enter the image from features that are outside the field of view.

We note that information in single-sideband images is “relocalized” by the usual CTF correction (Downing and Glaeser, 2008), even at spatial frequencies at which Thon rings are no longer visible. Since delocalized single-sideband information is recoverable to the same extent as when its Friedel mate is still in the field of view, it can be advantageous to use higher defocus values to record images (Russo and Henderson, 2018b). This remains true even for defocus values so high that the Thon-ring modulations become greatly reduced.

The second category of envelope effect is mentioned only for completeness, and we do not consider it further below. It results from delocalization of high-resolution information by a distance equal to more than half the field of view, a situation that did not occur in the experiments presented here. When this happens, as it might if the field of view is reduced by using much higher magnification to record images, both of the high-resolution Fourier components may begin to fall outside the area recorded by the camera. As a result, the average value of the background-subtracted power spectrum from features inside the field of view is expected to fall to zero, as does the modulation depth of the CTF. This effect happens first for particles near the center of the field of view. It is self-evident that images must be recorded with a smaller amount of defocus than that which produces this second type of envelope function. As before, information from single sidebands arising from features outside the field of view will contribute to the power spectrum.

## MATERIALS AND METHODS

### Specimen

The specimen used to obtain the data presented here consisted of a Quantifoil Cu R 2/2 grid (www.quantifoil.com), which was made hydrophilic by treatment with a glow discharge, to which 10 nm gold nanoparticles (www.cellbiology-utrecht), diluted 1:10 in water, were then applied. The thickness of the holey, amorphous-carbon film, estimated by comparing the Thon-ring power at ~8 Å with that from a calibrated carbon film of known thickness, was ~350 Å but had thicker and thinner regions. A typical area of the specimen is shown in Fig. 1A, in which 17 gold nanoparticles are visible in an area that measures about 0.2 x 0.2 μm. Each nanogold particle contains crystalline domains that are typically about half the diameter (5 nm) of each particle. Images obtained with other specimens were also analyzed in the course of work reported here, including one in which gold particles were deposited on a thin carbon film that was laid over the holes of a Quantifoil grid. Although the same Thon-ring behavior was seen with these, none had a distribution of gold particles as uniform as that presented here.

**Figure 1.**
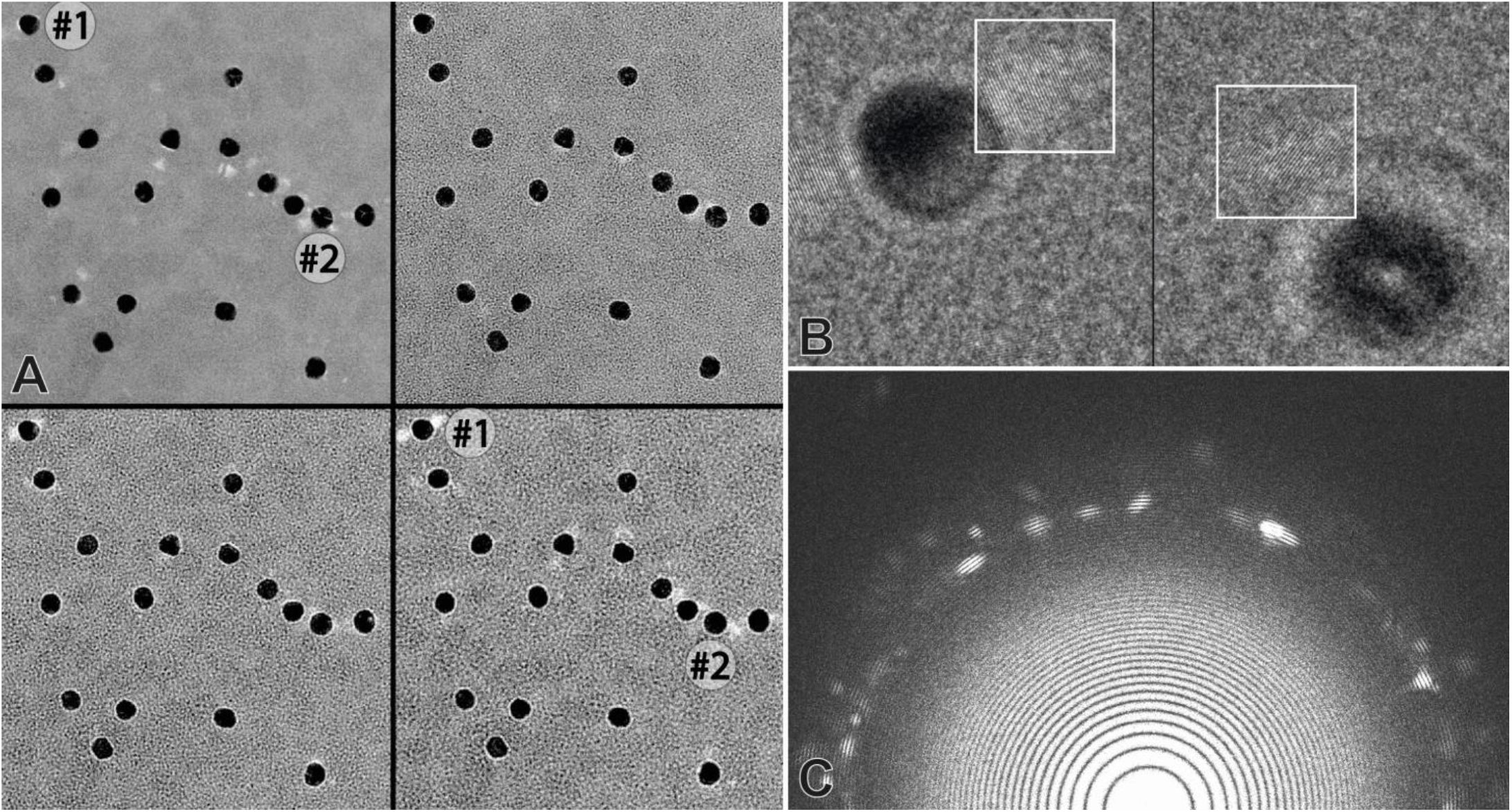
Examples showing the focus-dependent delocalization of information carried by interference between scattered electrons and the unscattered, reference wave. (A) Low-magnification view of the 4 images (gold-01 to gold-04) taken with the least amount of defocus, namely 0.02, 0.43, 0.87 and 1.3 μm. The strongest diffraction from an individual gold nanoparticle can be seen in the top left particle, labelled #1, of the last (1.3 μm defocus) image shown. This is the same particle shown at higher magnification in the left panel of Fig. 1B. Particle #2, whose fringes are shown in the right panel of Fig. 1B, is also labelled. (B) Subregions of two images showing the gold (111) fringes at 2.35 Å spacing. These images are for particle #1 from gold-04 image with 1.3 μm defocus, and for particle #2 from gold-20 image with 8 μm defocus. The region shown in the right panel is region B in Fig.3, showing that the gold-lattice fringes from particle #2 are separated by 700 Å from the image of particle #2 itself. (C) Fourier transform of the image recorded at a defocus value of 1.3 μm (gold-04), showing the Thon-ring sampling of the Au (111) spots.

### Imaging conditions

Data analyzed in this paper comprise a series of 20 images, in which the focal increment between successive images was about 425 nm. Summarizing briefly, all images were recorded on a FEI (now Thermo Fisher) Krios G1 microscope, using a Gatan Bioquantum energy filter and a Gatan K2XP camera. The full conditions used to record images are given in Table 1. Images were recorded as “movies” consisting of 40 frames each, for which the exposure time per frame was 0.5 s. Although the total exposure time per image, and the corresponding electron exposure per image, was about 4 times greater than that normally used in single-particle cryo-EM, the spatial coherence of the illumination used to record the images remained fully comparable.

**Table 1:** Technical details for 20 images in the defocus series of nanogold particles on carbon

| Technical details |  |
| --- | --- |
| Microscope | Thermo Fisher Krios G1 |
| Operating energy | 300 keV |
| Cs / Cc | 2.7 mm / 2.7 mm |
| Estimated beam tilt | < 0.1 mrad in first image ($\Delta F = 0$) |
| Energy filter and window | Gatan BioQuantum, 20 eV |
| Camera | Gatan K2XP |
| Exposure type | Counting mode / 40 frames aligned on-the-fly in SerialEM |
| Pixel size / magnification : calibrated / nominal | 0.508 Å / 118,000x / 75,000x (first image) |
| Exposure time per image | 20 s |
| Stage drift rate | 0.2 Å/s |
| Electron dose per image | 154 el/Å<sup>2</sup> = 40 el/pixel |
| Defocus range for series of 20 images | 0.02 µm – 8.3 µm, in 19 steps of $\Delta F = 4340$ Å |
| Astigmatism | < 70 Å |
| Linear magnification anisotropy | +0.09 % at angle 53°, -0.09 % at 143° |
| Carbon film thickness | ~350 Å |
| Nanogold particle diameter | 100 Å |
| Nanogold crystal domain size | 50 Å |
| Location of gold particles | On lower side of carbon film in microscope |
| Gold 111 spacing used for calibration | 2.35 Å |
| Specimen temperature | -186°C |
| Magnification change from image 1 to 20 | + 0.8 % (equivalent 0.18 mrad convergence) |

The microscope alignment was checked with AutoCTF for stigmation and coma-free alignment. We estimated any residual astigmatism to be less than 70 Å, and the remaining beam tilt to be less than 0.1 mrad. The camera pixel size at the specimen was 0.508 Å, calibrated from the Au (111) reflection on the first (zero defocus) image. The magnification on the 20^th^ image, taken with an additional 8 μm underfocus, was 0.8% higher, and this was accounted for during data processing.

### Systematic corrections were applied before calculating 1-D CTF curves

Images were first floated and padded to slightly more than twice their initial size, as described more fully in the next section, before being subjected to Fourier transformation (FT). This was done in order to have enough sampling of the FT to resolve what can become very closely-spaced Thon rings at larger-than-usual amounts of defocus. In particular, a perhaps-subtle effect occurs when the separation between two Friedel-related, delocalized contributions to the image becomes equal to half the width of the image, or more. Briefly, the period of the resulting Thon-ring modulation at that spatial frequency becomes less than two pixels in the FT unless the image is padded.

A preliminary calculation of the circularly averaged FT was carried out for every image. Examination of these plots showed that the circularly averaged Thon-ring modulation at the gold (2.35 Å) spacing faded out at about 5 μm defocus and then reappeared at higher values of defocus. This suggested the presence of a small amount of linear magnification anisotropy, which made the Thon rings very slightly elliptical. The Thon-ring spacing at 5 μm defocus was then used to give an initial estimate of 0.17% linear-magnification anisotropy.

In a second step, which is described more fully in Appendix 1 of the Supplemental Material, this initial estimate was used to refine the magnitude and direction of the linear magnification anisotropy (two parameters). For our particular data set, the final value proved to be 0.17% anisotropy, close to the initial estimate, and at an angle of 53°. The correction applied consisted of an increase in the radius for the power-spectrum binning by 0.09% at 143° and a corresponding decrease at 53°, using the function cos(2θ), where θ is the angle between the defined anisotropy direction and the vector direction to each pixel. The presence and amount of linear magnification anisotropy may depend upon the values of stigmator currents set during Supervisor alignment. Making this correction has no effect on the appearance of the circularly averaged Thon rings at small values of defocus (up to 2 μm), but it significantly improves their visibility at defocus values higher than 4 μm. It is important to note that the Thon rings are present in all directions, regardless of the amount of defocus, but the slight ellipticity that existed, prior to the anisotropy correction, damped the modulation in the circular average at high values of defocus.

### Calculation of 1-D CTF curves

The 1-D power spectra were calculated using single-purpose scripts and new code added to the package of MRC software (Crowther et al., 1996) as well as adapted local routines. Briefly, the images were padded out to 8192×8192 pixels from the original size of 3708×3838 pixels, and the average values of the power spectra were calculated in annular zones of 1 reciprocal-space pixel. The padded images were floated in order to avoid production of streaks through the origin of the Fourier transform, due to an abrupt discontinuity between the average intensity of the original image and that of the surrounding, padded area. The Nyquist resolution is 1.016 Å, at the 4096^th^, edge pixel in the FFT, and the gold 111 Fourier component is at just less than half the Nyquist frequency.

Due to the padding, the width of the annular zones near the gold 2.35 Å spacing was reduced to less than 0.33 times the width of the Thon rings, even for the 20^th^ image in the series, recorded with the greatest amount (8 μm) of defocus. There was no further improvement in the visual clarity or the amplitude of the Thon rings achieved by using greater amounts of padding, for example out to 16384×16384.

The effect of astigmatism on Thon-ring modulation in a radial power spectrum increases at high resolution but is independent of the defocus. Since the amount of astigmatism was estimated to be less than 70 Å, the effect of astigmatism on Thon-ring modulation at the gold spacing of 2.35 Å was expected to be less than 1/6^th^ of the Thon-ring spacing.

### Measurement of the CTF-modulation depth

The following analysis was performed in order to measure fading of the Thon rings in a particular frequency band, Smooth curves were first fitted to the local maxima and minima, respectively, of the Thon rings for each of the power spectra. For the 9 Å and 7 Å bands, whose power is from the carbon alone, the mean value minus the minimum value, at a particular defocus, was then taken as a measure of the intensity of the Thon rings in that band. These intensity values were then normalized by dividing by the value obtained for the first in the series of spectra, as well as fitting an offset to account for the first measurement in the series not being at 0 defocus. Low-defocus spectra were not included in the analysis if they lacked a full period of oscillation in the band of interest. The 9 Å band included data from 10.4 Å to 8.3 Å, and the 7 Å band was from 8.3 Å to 5.9 Å. For the 2.3 Å band, which is dominated by scattering from gold 111 reflections, the maximum minus the minimum in the band between 2.38 and 2.33 Å was taken as the intensity, and it was again normalized in the same way.

## RESULTS

The field of view presented in Fig. 1A shows there are 17 gold nanoparticles in the particular series of 20 images analyzed in this paper. For ease of reference, individual members of this focal series are referred to as “gold-01” through “gold-20”, respectively, beginning with the one that was recorded with the least amount of defocus. The degree to which the intensity of the delocalized information varies from one gold particle to the next, clearly evident in Fig. 1A, is expected to depend in a very sensitive way upon the orientation of each particle with respect to the incident electron beam.

The two gold nanoparticles labelled #1 and #2 are discussed in more detail. The strongest lattice images are seen from particle #1 in the bottom right panel of Fig. 1A, which has a defocus value of 1.3 μm. This particle, at the same defocus, is shown at higher magnification in the left panel of Fig. 1B, whereas the gold fringes from particle #2 from the 8 μm defocus image (gold-20) are shown in the right panel. The integrated power at 2.35 Å for particle #2 is comparable to that of particle #1 (67% of intensity), even though the image is much more highly defocused (8 μm as opposed to just 1.3 μm). The FT of the entire 1.3 μm defocus image gold-04 is shown in Fig. 1C, where Thon-ring sampling of the gold diffraction spots can be clearly seen, many at 2.35 Å and 2.04 Å, and, very faintly, some even out as far as 1.4 Å.

As is shown in Fig. 2, the circularly averaged power spectra exhibit defocus-dependent oscillations (Thon rings) extending to a resolution beyond 0.2 nm, even when the images are recorded at a defocus value of 4 μm or more. The power spectra also contain a broad peak at a resolution of ~0.23 nm, due to Bragg diffraction from the nanocrystals. This relatively broad peak provides an internal size-standard, which was used to calibrate the spatial-frequency axis. The width of the broad peak is ~1/50 Å^−1^, implying that the size of the crystalline domains within each gold nanoparticle is ~50Å. The amplitude of the Thon-rings that modulate this broad diffraction peak is much greater than it is in the Fourier transform on either side of the peak, an effect that is due to the structure factor for the nanocrystals being much greater than it is for the amorphous carbon film.

**Figure 2.**
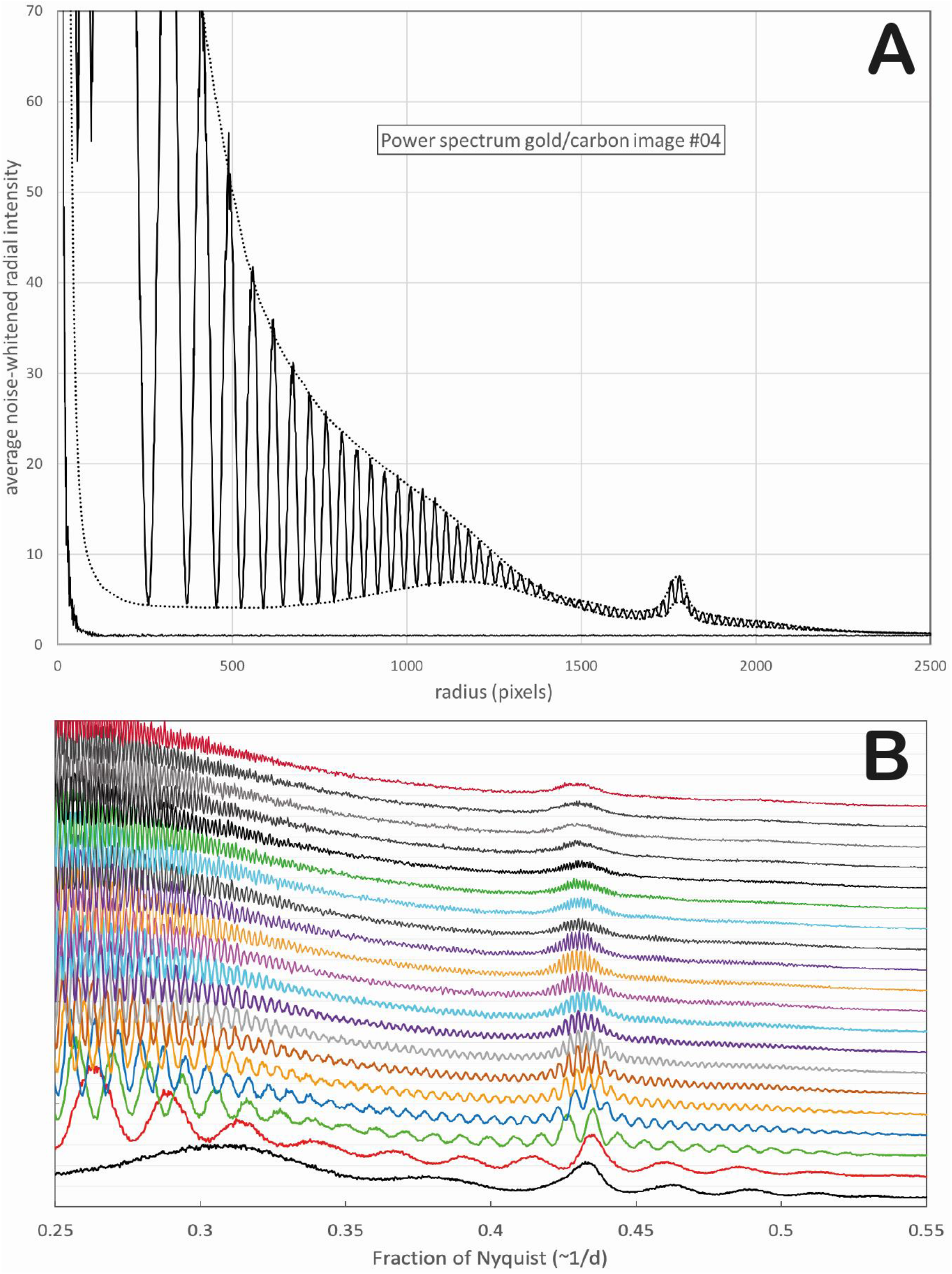
1-D power spectra of images recorded at various values of defocus. (A) The power spectrum for the image taken at 1.3 μm defocus, along with that of an image recorded without a specimen, are shown to illustrate the methodology of the data analysis presented in this paper. (B) Power spectra of the region between 0.25-0.55 Nyquist for all 20 images, offset from one another in order to see the gradual changes in the modulation of Thon rings that occur as the amount of defocus increases from 0.0 μm to 8.0 μm, in defocus steps of 4250 Å.

The power spectrum of the image recorded at a defocus value of 1.3 μm is shown in Fig. 2A. Dotted-line curves in this figure represent fits to the maxima and the minima, respectively of the circularly averaged Thon rings. Fig. 2A also includes a plot of the power spectrum of an image recorded without a specimen, which provides an example of the additive “shot” noise expected in images. The mean value of this noise spectrum can be regarded as the “zero baseline” for the power spectra of images recorded with a specimen.

We draw attention to the fact that the CTF minima do not touch the “zero” baseline, i.e. the expectation value of the power spectrum of additive noise. Instead, there is a gap between the power spectrum of noise and a curve fitted to the minima of the oscillations, as can be seen in the example shown in Fig. 2A. As is addressed further in the Discussion, this gap demonstrates that the image contains small, yet measurable contributions that are not accounted for by the weak-phase approximation.

The amplitude of the Thon-ring oscillations decreases progressively as the defocus value is increased from 2 μm to 6 μm, as is shown in Fig. 2B. Although it becomes increasingly difficult to see Thon rings over much of the power spectrum at higher values of defocus, they nevertheless remain clearly detectable within the broad, 111 diffraction ring of the gold nanoparticles – where the signal is much higher, even at defocus values well above 6 μm. The cause of the decreasing amplitude of the Thon rings is addressed in the Discussion, below.

## DISCUSSION

### The observed, focus-dependent reduction in Thon-ring amplitude is not due to a spatial-coherence envelope function

As mentioned in the Introduction, the historical concern about the spatial-coherence envelope function (Wade and Frank, 1977) still discourages many cryo-EM users from recording images at high defocus. Indeed, the supposed limitation due to such an envelope function was cited as a reason for using ptychography rather than bright-field imaging (Zhou et al., 2020). Nevertheless, the high brightness of Schottky field-emission guns makes it possible for the illumination semi-angle to be as low as 7 μrad under low-dose conditions (Börrnert et al., 2018), which is much smaller than for thermionic sources, or that assumed in the simulations in (Zhou et al., 2020). Thus, under the low-dose conditions used for cryo-EM, the spatial-coherence envelope function is not expected to limit information transfer out to a resolution of 2 Å, for example, until the defocus value is increased to about 4 μm or more [see Figure 6 in (Börrnert et al., 2018)]. Experiments showing that nearly all of the signal in delocalized lattice images of gold nanocrystals is retained (Russo and Henderson, 2018a) have demonstrated that the spatial-coherence envelope function is no longer a concern at defocus values as high as 4 μm or more, and similar results are presented here, confirming those results.

We note that spatial coherence (for the illumination conditions used here) did not prevent lattice images at 8μm defocus from being recorded at a resolution of 1.44 Å, corresponding to the 220 Bragg spacing of gold. Although the temporal-coherence envelope is expected to attenuate the signal significantly at that resolution, such fringes were nevertheless seen in many areas of the images recorded at defocus values as high as 6 μm. This leads us to suspect that the illumination semi-angle may have actually been somewhat less than the value of 7 μrad cited above. Indeed, we calculate that it must be in the range 1-2 μrad, based on our measured dose rate having been 7.7 e/Å^2^-s and an estimated value of ~5×10^7^ A/m^2^/sr/V for the SFEG gun brightness, which is given in the Titan Condenser Manual. Under lower dose-rate conditions of about 2 e/Å^2^-s, often used in cryo-EM, the illumination semi-angle might even be less than 1 μrad. As we now report, there nevertheless is a progressive, defocus-dependent reduction in the amplitude of Thon rings, which becomes especially evident when the defocus value is increased beyond 5 or 6 μm. In spite of this, the signal continues to be retained very well in the delocalized lattice images of gold nanocrystals. This is not what is expected if the spatial-coherence envelope were responsible.

Even the diminishing amplitude of Thon-rings fails to behave in the way expected if the cause was limited spatial-coherence. For example, the values of local minima in the power spectrum gradually increase at the same time as the values of the maxima decrease, i.e. as the amount of defocus increases, which is not the behavior expected for a spatial-coherence envelope function. Instead, a more-weakly modulated signal (and eventually an unmodulated signal) is retained, whose power remains midway between the values of the local maxima and minima of the Thon rings that are seen at lower values of defocus. This progressive convergence to an intermediate value is also seen in the Thon-ring modulated, broad diffraction ring from the gold nanocrystals, at a resolution of ~2.35 Å, as is seen in Fig. 2B.

We wanted to know what the correct explanation might be for the focus-dependent reduction in amplitude of the Thon rings, since the spatial-coherence envelope is ruled out. As we discuss below, we found that the focus-dependent delocalization of information appears to account what we found experimentally, while various forms of non-isoplanacity make only negligible contributions.

As an aside, we also note that the local minima in the power spectrum of both the carbon film and the gold nanoparticles are above the “zero baseline” value that corresponds to the power spectrum of shot noise (see Fig. 2 A, for example). In effect, the background-subtracted power spectrum does not go to zero at the zeros of the CTF. It thus is clear that the weak-phase, weak-amplitude object approximation does not account for all of the power in the Fourier transforms of the images like those shown in Fig. 2. Some of the discrepancy might be due to quadratic terms (dark-field images), which are usually ignored in theoretical expressions of the intensity of images of weak objects. Little of it can be attributed to the image intensity produced by inelastically scattered electrons, however, since an energy filter was used to record the images presented here. Finally, we note that there is an unexpected, broad bump in the apparent baseline that corresponds to the local minima in the Thon-ring oscillations, peaking at a resolution of about 3.5 Å. We identify one factor, in Appendix 3 of the Supplemental Material, that contributes to formation of the bump. None of these features bear on the question of whether spatial-coherence is a limiting factor out to a resolution of ~2Å, so we do not discuss them further here.

### Delocalization accounts for the observed reduction in Thon-ring amplitude

The observed, focus-dependent reduction in the modulation depth of Thon rings is well accounted for by a wavelet-inspired model, in which sinusoidal packets of structural information are progressively delocalized outside of the field of view of the camera. As this is not a commonly used model to describe delocalization of information in the cryo-EM context, we begin this section of the Discussion with a brief description of the model. We then apply the model to the two categories of defocus-dependent behavior of the CTF that are referred to in the Introduction.

#### Wavelet representation provides a convenient description of defocus-dependent delocalization

As background, we here consider the spatially varying phase of the electron wavefront immediately after it has passed through a small particle, i.e. the exit wave, to be represented as a superposition of wavelets, i.e. sinusoidal components of the projection of the structure, each of which is confined to the boundaries of the particle. Such a wavelet decomposition provides a natural description when the particle is a nanocrystal, in which case the period of oscillation corresponds to a Bragg spacing of the crystal, and the compact support of the wavelet corresponds to the physical boundaries of the nanocrystal. The same picture is easily generalized to a non-crystalline particle, of course, and even to an imagined nano-region of a continuous, amorphous-carbon film.

As the transmitted electron wave propagates beyond the exit face of the particle, every such wavelet divides into two, delocalized patches, which we will refer to below as “half-wavelets”. After propagating far from the exit face, each such patch of the electron wave function eventually becomes a spot in an electron diffraction pattern. For a weak-phase object, these half-wavelets each have half of the original amplitude of the full wavelet, and equal and opposite spatial frequencies.

In the plane of a defocused image, i.e. much closer to the particle, the centers of these patches are shifted (delocalized) to either side of the center of the particle by an amount that is equal to the gradient of the wave-aberration function of the imaging lens, evaluated at the frequency of oscillation of the wavelet. Considering only defocus, the displacement, *α*, is given by

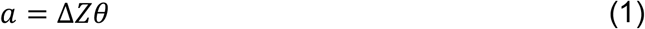

where Δ*Z* is the defocus of the image, *θ = λ/d* is scattering angle, in which *λ* is the electron wavelength and *d* is the Bragg spacing.

Once the amount by which patches are shifted becomes greater than the size of the particle, these delocalized half-wavelets produce separate, single-sideband contributions to the image intensity. Furthermore, although the amplitude of oscillations within single-sideband components is independent of focus – see, for example, section 3.11 of (Glaeser et al., 2007), the Fourier transform of the whole image continues to be modulated by the CTF. CTF modulation of the Fourier transform is conveniently attributed, in this case, to interference between the contributions from the two patches, which can be either constructive or destructive, depending upon the distance by which the patches are separated, which is *2a* (see Equation 1).

Importantly, this model assumes that constructive and destructive interference occurs between the respective, delocalized half-wavelets and the reference wave, i.e. the unscattered beam, during formation of the image, thereby producing lattice images outside the outline of the particle itself. The persistence of lattice images within the boundary of a half-wavelet is thus proof that the coherence diameter is equal to or larger than twice the distance by which the half-wavelet is displaced from the particle.

#### CTF modulation disappears when only one component of delocalized information is recorded

To summarize the preceding section, as long as both of the delocalized patches remain within the field of view, the Thon-ring modulation of the Fourier transform of the images varies with defocus in the usual way. This no longer remains true, however, if one of the patches is delocalized beyond the limited field recorded by the camera. At that point the remaining patch still contributes a share to the power spectrum, but it no longer contributes to the CTF oscillations.

Particles that lie closest to the edge of the field of view are obviously the first to have one of their half-wavelets become delocalized outside of the recorded image. As the amount of defocus increases further, particles located closer and closer to the center of the field of view will experience the same effect, i.e. only one of their half-wavelets will remain within the field of view. An example illustrating this point is shown in Fig. 3. As a result, the fraction of the field of view for which both half-wavelets are retained continues to decrease as the amount of defocus increases. As is derived in Appendix 2 of the Supplemental Material, the resulting area that still contributes to Thon-ring modulation decreases almost linearly with how far the half-wavelets are displaced. The rate at which the modulation depth decreases also depends upon the spatial frequency of interest, since the delocalization itself increases linearly with spatial frequency. As the examples in Fig. 4 demonstrate, this wavelet model describes our experimental data. We note that this behavior is different to what is expected if the focus-dependent fading of the Thon rings were caused by a spatial-coherence envelope function.

**Figure 3.**
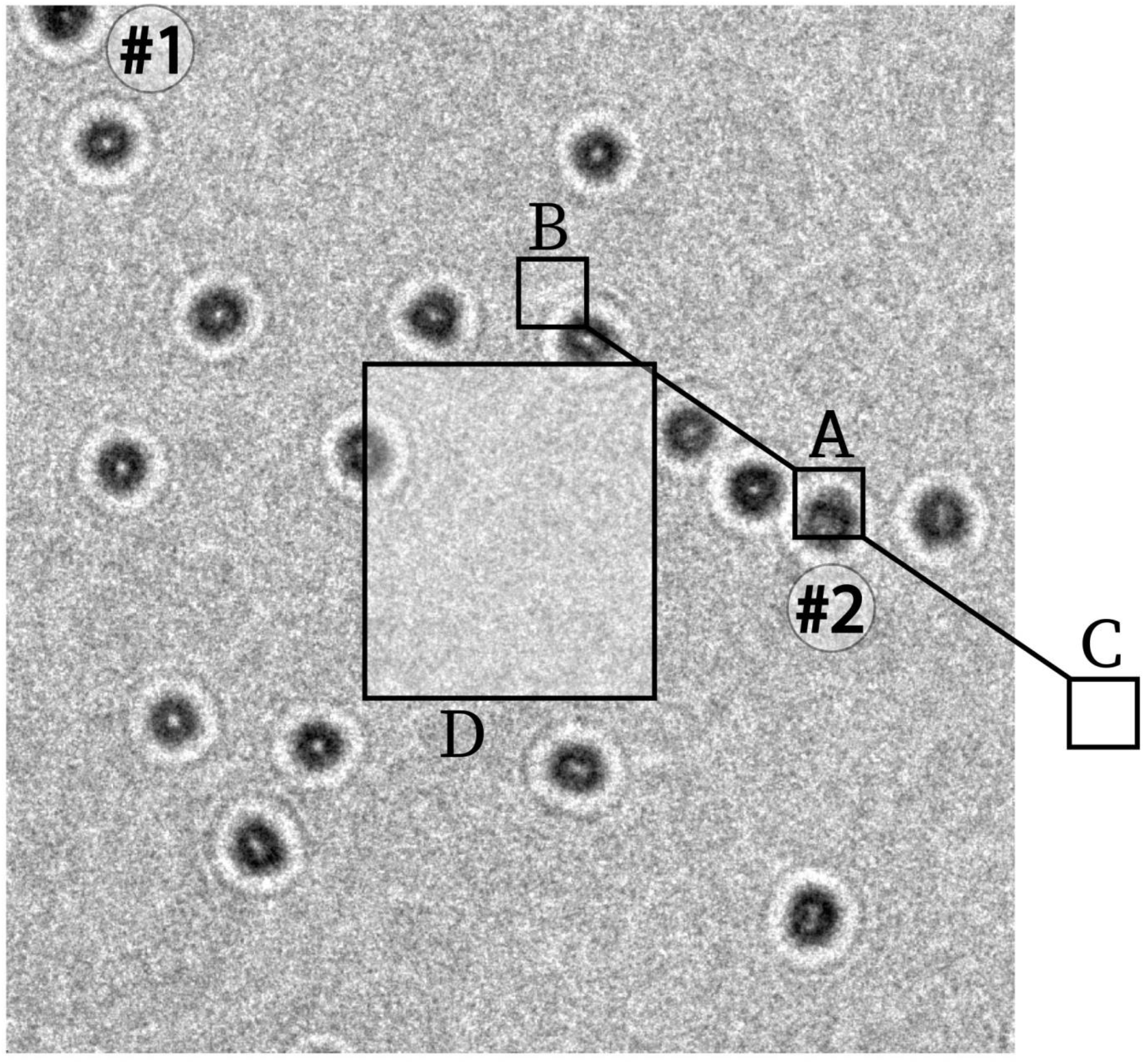
Image “gold-20”, recorded at 8 μm defocus, showing how delocalization of one of the half-wavelets corresponding to the Au (111) lattice planes is moved outside the field of view of the camera. Gold nanoparticle #2 is labelled, with the diffracting crystallite labelled A, and the positions of 2.35 Å lattice fringe patches labelled B and C. Area C is outside the field of view, so this half-wavelet is not observed and it does not contribute to the power spectrum. Area B is shown at higher magnification in the right panel of Fig. 1B. The area outlined in the center would be the only region able to produce Thon-ring modulations at the 2.35 Å gold-lattice spacing in all directions, but, within this field of view, there is only about half of one of the 17 gold particles that is located within the outlined area. As a result, the power spectrum of this image shows no Thon-ring contribution to the power spectrum of such nanoparticles.

**Figure 4.**
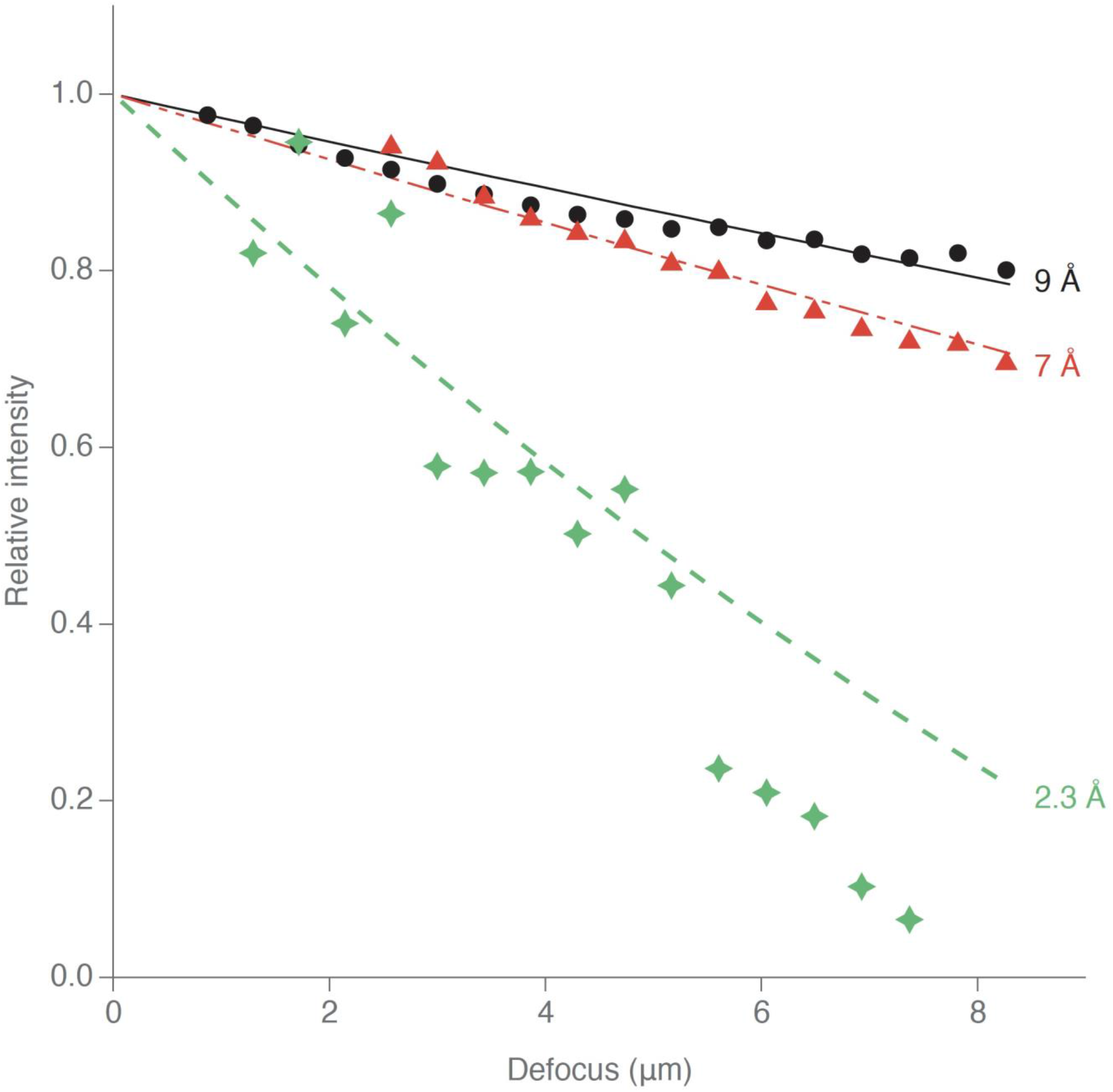
Plots showing how the Thon-ring modulation depths, in different resolution zones, decrease as the defocus is increased. Theoretical curves show the fraction of the area of the field-of-view for which both wavelets remain in the recorded image, whereas circles (9 Å), triangles (7 Å), and diamonds (2.3 Å) represent the experimental data shown in Figure 2B.

The value of defocus at which the modulation depth finally reaches zero depends both upon the spatial frequency of a Fourier component of the specimen and upon the size of the camera (i.e. the pixel size and the number of pixels), referred back to the specimen. According to Equation 1, a half-wavelet with a period of 2.3 Å is expected to be displaced from the original particle by a distance of ~870 Å when the image is defocused by 10 μm. This displacement is almost half of the field of view under the experimental conditions used here. Thus, at a defocus of 10 μm, only a tiny fraction of the area of the image still remains in which particles contribute both of their delocalized half-wavelets to the image. An analytical expression for this fraction of the area is given in Eq. S6 of Appendix 2 of the Supplemental Material. Numerical simulations of Thon ring fading are presented in Appendix 3 of the Supplemental Material, and a table of calculated displacements is presented in Appendix 4 of the Supplemental Material, for electron energies of 100 keV and 300 keV, respectively.

In practice, the highest defocus used here was 8 μm, so even here a small fraction of the image could in principle give rise to gold Thon-ring sampling. Unfortunately, as seen in Fig. 4, there is only half of one gold nanoparticle in that region and it may not be in a diffracting orientation. We therefore expect there to be nearly zero Thon-ring modulation of the gold diffraction in the final 8 μm defocus image in the series, as observed.

The discussion above assumes, of course, that the displaced half wavelets remain coherent with the unscattered-electron, reference wave. This is a good approximation for 300 keV electrons under low dose-rate conditions, as indicated by the fading experiment in (Russo and Henderson 2018a) and by the illumination semi-angle reported by (Börrnert et al., 2018).

#### The average-signal level is reduced by half when only one component of delocalized information is recorded

Let *F* be the amplitude of a scattered electron beam (and of its Friedel mate), also referred to as the magnitude of its structure factor. In the absence of a spatial-coherence envelope function or other losses, the amplitude of the Fourier transform of a double sideband image is proportional to 2F sin γ(θ), where *γ*(*θ*) is the wave aberration due to defocus and spherical aberration given by

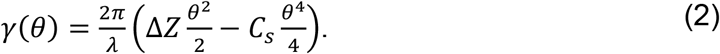

Under the same conditions, the amplitude of a single-sideband image is proportional to *F*, independent of the value of *γ*. The signal power in the Fourier transform of a single-sideband image is thus proportional to *F^2^*, which is one quarter the peak power, and one half the average power, in the Fourier transform of a double-sideband image.

In the experimental results shown in Fig. 2B, however, it is not apparent that the average power of the Fourier transform is reduced as the CTF modulation depth decreases. The reason that the average power remains constant is that “invading” half-wavelet patches, which originate from particles lying outside the field of view, are delocalized into the field of view. This only happens if sufficient area outside of the field of view is illuminated by the electron beam.

### Off-axis aberrations do not currently contribute significantly to the anomalous envelope effects

For completeness, we note that anything that causes the CTF within one sub-region of a recorded image to differ from that in another sub-region could, in principle, also produce an anomalous envelope function. Were that to happen, the image would no longer be isoplanatic, although smaller regions within the image might still be approximated as being isoplanatic. An isoplanatic patch, as defined in section 2.2 of (Goodman, 1968) or section 9.5 of (Born and Wolf, 1997), is an area within which the image of a point, although possibly distorted, is everywhere the same to within the resolution of interest.

Effects that may limit the size of the isoplanatic patch, and which are not compensated by making the systematic corrections described in this paper, include off-axis coma (Glaeser et al., 2011), some types of magnification distortion including that in the energy filter lens (Gubbens et al., 2010) or field aberrations encountered when stitching images (Kaynig et al., 2010), and variations in Z-height of different areas of what is otherwise a thin sample. The agreement shown in Fig. 4 between the experimental data points and the results expected because of delocalization of half-wavelets, indicates that delocalization alone can account for the focus-dependent loss. It thus appears that the images obtained here are isoplanatic to a resolution of 2 Å.

## SUMMARY AND CONCLUSIONS

The amount by which the amplitudes of Thon-ring modulations decrease at high values of defocus is investigated under conditions that are currently in use in single-particle cryo-EM. The results demonstrate that defocus values as high as 4 μm result in little loss of information about the specimen, up to a resolution of about 2 Å. Contrary to common beliefs, there is no significant disadvantage to record images with defocus values well above the usual practice, in order to increase the signal-to noise ratio at low resolution, if doing so improves the ability to extract and process images of individual particles.

Although the loss of information does become significant at defocus values of 6 μm or more, even this loss is not caused by limited spatial coherence, as is often claimed. Instead, we demonstrate that it is caused by the delocalization of information outside the field of view of the camera. In the future, it can be expected that data will be collected with pixel sizes smaller than 0.5 Å (referred to the specimen), in order to achieve resolutions that are twice, or more, than investigated here. In this case, delocalization will play a greater role, and corresponding care will have to be taken to record images within a correspondingly low, but still useful value of defocus.

## ACKNOWLEDGEMENTS

This work was completed in part with the support of NIH grant GM126011, MRC grants MC-U105184322 and MC-UP-120117, and the Structural and Computational Biology unit, European Molecular Biology Laboratory, Heidelberg.

## SUPPLEMENTARY MATERIAL

### Appendix 1: Using CTF fitting to determine anisotropic magnification

There are many ways to determine linear magnification anisotropy (Afanasayev et al, 2017; Grant & Grigorieff, 2015; Yu et al, 2015; Zhao et al, 2015), but if a defocus series is available, as in this work, then it is convenient to make use of established programs such as CTFFIND3 or CTFFIND4 (Rohou & Grigorieff, 2015) to measure how the apparent astigmatism varies with defocus, then to fit the anisotropy magnification parameters to explain the changes in apparent defocus and astigmatism. Small amounts of astigmatism and small amounts of anisotropic linear magnification have the same effect, which is to stretch the image and the Fourier transform in one direction. Of course, the two stretches are normally in different directions. They can both be characterized as a variation dependent on cos(2*φ*), where the angle *φ* is measured from two different directions, namely the true astigmatism direction and the true linear magnification anisotropy direction. Ignoring astigmatism, the magnification at an angle *φ* in the presence of anisotropic magnification can be written as

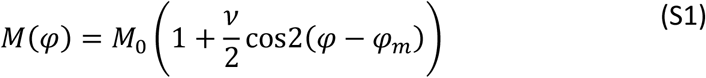

where *v* is the magnitude of anisotropy and *φ_m_* is the direction of the linear magnification anisotropy. As a result of the linear magnification anisotropy the sampling of the specimen seen by a detector will be direction-dependent. In the FFT of an image from a square detector of size *N* with pixel size *p*, the true scattering angle at pixel (i,j) is

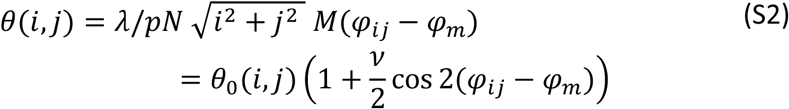

where *φ_ij_* is the angle relative to the pixel array of the ij-th pixel from the origin and *θ*_0_(*i,j*) is the expected scattering angle with isotropic magnification. The wave aberration due to defocus depends on the true scattering angle and, as seen in the ij-th pixel of the FFT, is

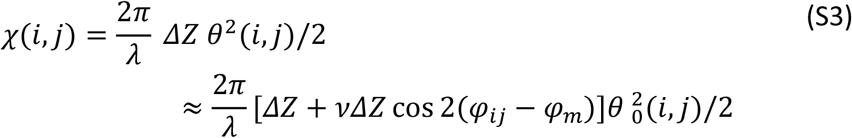

which is identical to that seen with isotropic magnification, defocus of *ΔZ*, and astigmatism of 2*vΔZ*. The linear isotropic magnification can therefore be determined by simply plotting the measured astigmatism as a function of defocus and extracting the slope. In the main paper, CTFFIND3 and CTFFIND4 were used to find the defocus and astigmatism, and a program astiganisotropy.py, which is available on request, was used to extract the linear anisotropic magnification and its direction. The output of astiganisotropy.py is shown in Fig. S1.

There will also be true astigmatism, but, by careful stigmation at low defocus, this can be minimized. As shown in Fig S1, with increasing defocus this contribution is quickly swamped by the anisotropy contribution. Note that true astigmatism does not change with defocus. We confirmed this by comparing two images recorded with the same amount of defocus but with the specimen at z-height locations that were 8 μm apart (i.e. with different lens excitation values).

**Figure S1.**
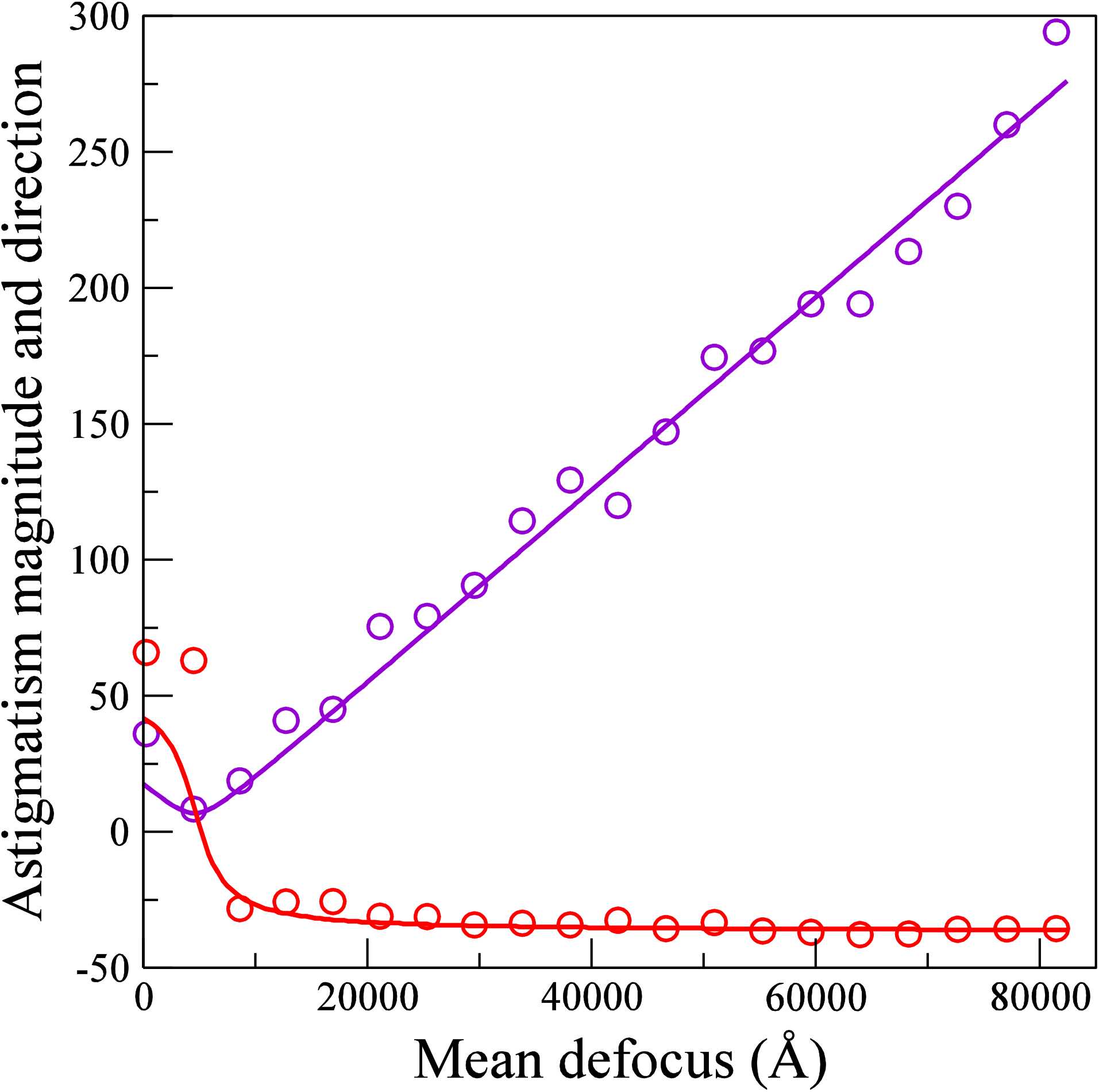
The values of the apparent astigmatism magnitude (purple) and direction (red) determined experimentally from the 20 images (symbols) in the defocus series using CTFFIND3 and CTFFIND4 are plotted together with the predicted values from astiganisotropy.py (continuous line). A linear magnification anisotropy of 0.18±0.01 % in the direction −37° and a residual astigmatism of 18 ±20 Å at 42° were found.

## Appendix 2: Derivation of Thon Ring fading with defocus for a finite size detector

Thon rings arise from the interference between the unscattered-electron wavefront and the phase-shifted, scattered-electron beams from a particular region of a sample in an image. In the case of the gold nanocrystals, it is convenient to consider the Bragg-diffracted beams, and the equivalent description is easily generalized. The phase shift of the Bragg beams, or sidebands, is spatial frequency dependent and to lowest order arises from the difference in path length due to defocus and spherical aberration of the lens. The generalization of Eq. 1 to include spherical aberration is simply

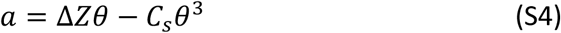

where a is the sideband displacement and C_s_ is the spherical aberration coefficient. The spherical aberration term adds a defocus independent offset which is typically only important at very low defocus or high resolution. Apart from the contribution of spherical aberration, the displacement will be independent of electron wavelength if the product of defocus and wavelength is held constant. Note that in the low defocus image of Fig. 1A, the strongest visible diffraction spots come from the [222] and [311] Bragg planes at ≈1.2 Å. With the convention of Eq. S4, these spots have a displacement of −114 Å. With increasing defocus these diffraction spots move towards, through and then away from the unscattered image. For reference, tables showing the expected values for the side band displacement for 100 and 300 keV electrons at various spatial frequencies and values of defocus are given in Appendix 4.

**Figure S2.**
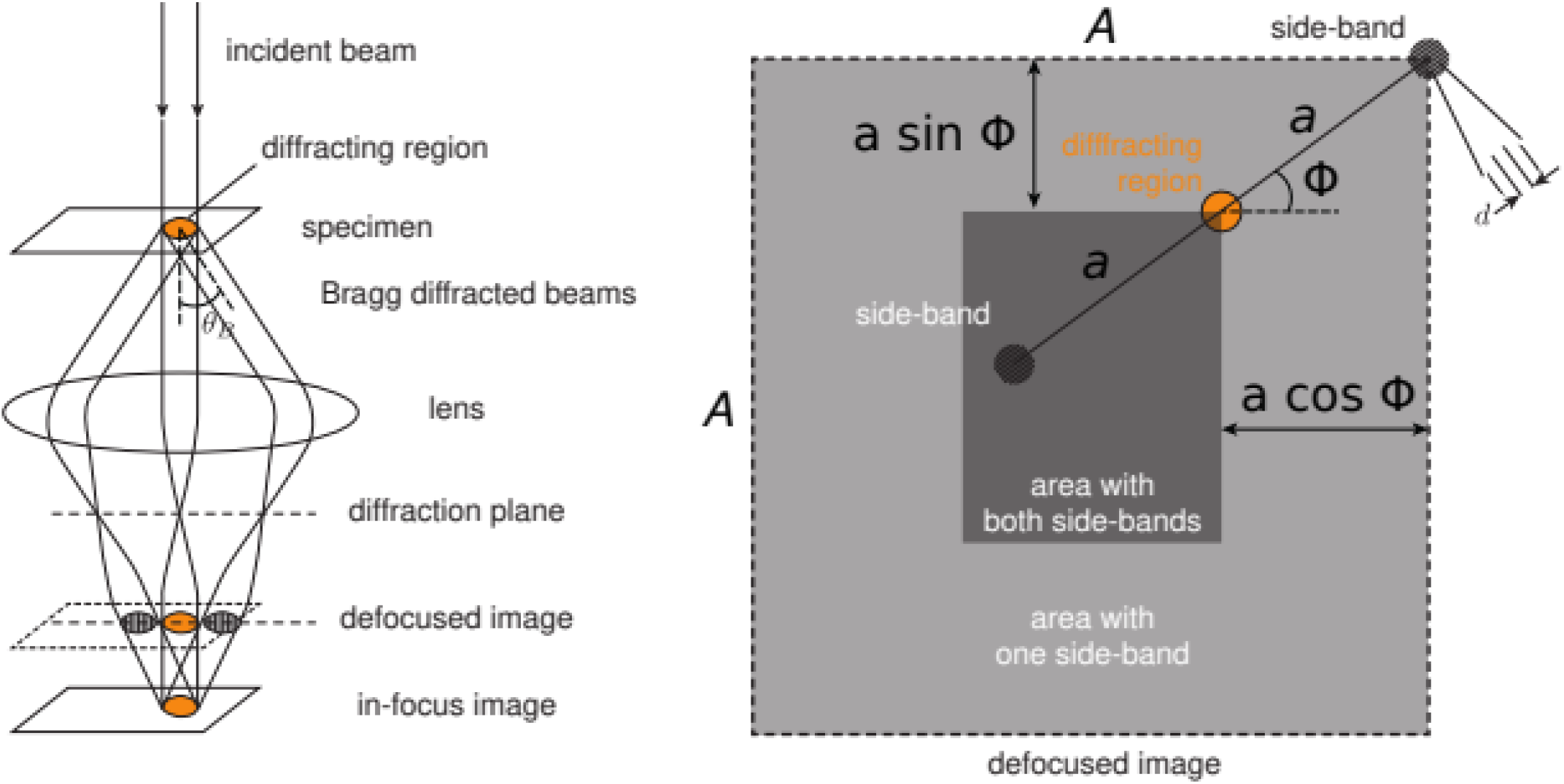
Illustration of the origin of the defocus dependent damping of the Thon ring modulation with a finite size detector. A ray diagram for the formation of a defocused image from a sample with a Bragg plane separation of d showing the origin of the sideband displacement is shown on the left. The part of the image (darker area) that contains both sidebands of a diffracted beam with a sideband displacement a, at angle *ϕ* to the detector, is shown on the right. With increasing defocus, the magnitude of the displacement increases and the fraction of the image that contains both sidebands is reduced.

The formation of a defocused image from a single Bragg plane is illustrated in the left-hand side of Fig. S2. In the absence of spherical aberration, the lens focuses back the Bragg diffracted beams to merge with the direct beam at the in-focus image plane. With defocus and spherical aberration, the diffracted beams are displaced from the direct beam by a distance *a* given by Eq. S4. Real detectors are of finite size and even though the low-resolution image of an object may be recorded on the detector, the delocalized diffracted beams may not be. This is illustrated in the right-hand side of Fig. S2 for the case where the sidebands are displaced by a distance *a*, at an angle *ϕ* to the detector array. Both sidebands will be recorded only if the unscattered beam falls in the indicated dark area in the center of the detector.

As noted earlier, if both sidebands are not present in the image, that part of the specimen will not contribute to the Thon ring modulation seen in the power spectra of the image. To develop an expression for how the Thon ring modulation is reduced as a result of delocalization, we consider a square detector of dimension *A*, and assume that there is uniform scattering throughout the specimen plane. The fraction of image that contributes both sidebands, as indicated in the right-hand side of Fig. S2, depends on the magnitude of the ratio of the sideband displacement to the detector dimension and the angle, *ϕ*, of the displacement relative to the detector array. If we define *α* = |*a*|/*A*, the fraction at angle *ϕ* that contributes both sidebands, *f*(*α, ϕ*), is given by

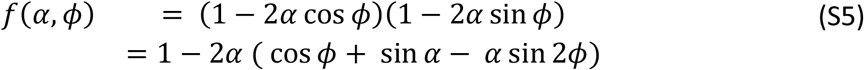

The fraction of the image that contributes both sidebands is found by averaging over *ϕ*. By symmetry the average need only be carried out over 0 to π/4. However, there are three regimes depending on the value of *α*. If *α* ≤ 1/2, the average is carried out over all *ϕ*. For 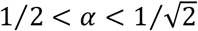, *f*(*α, ϕ*) is only nonzero for *ϕ* between cos^−1^1/2*α*. and π/4 Finally, for 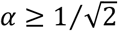, *f*(*α, ϕ*) = 0 for all *ϕ* because in this case at least one sideband is always scattered outside the area of the detector no matter where the unscattered image falls on the detector. With this in mind averaging over *ϕ* gives

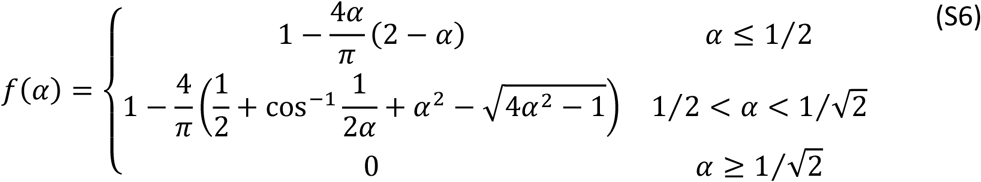

The behavior of *f*(*α*) is shown in Fig. S3

**Figure S3:**
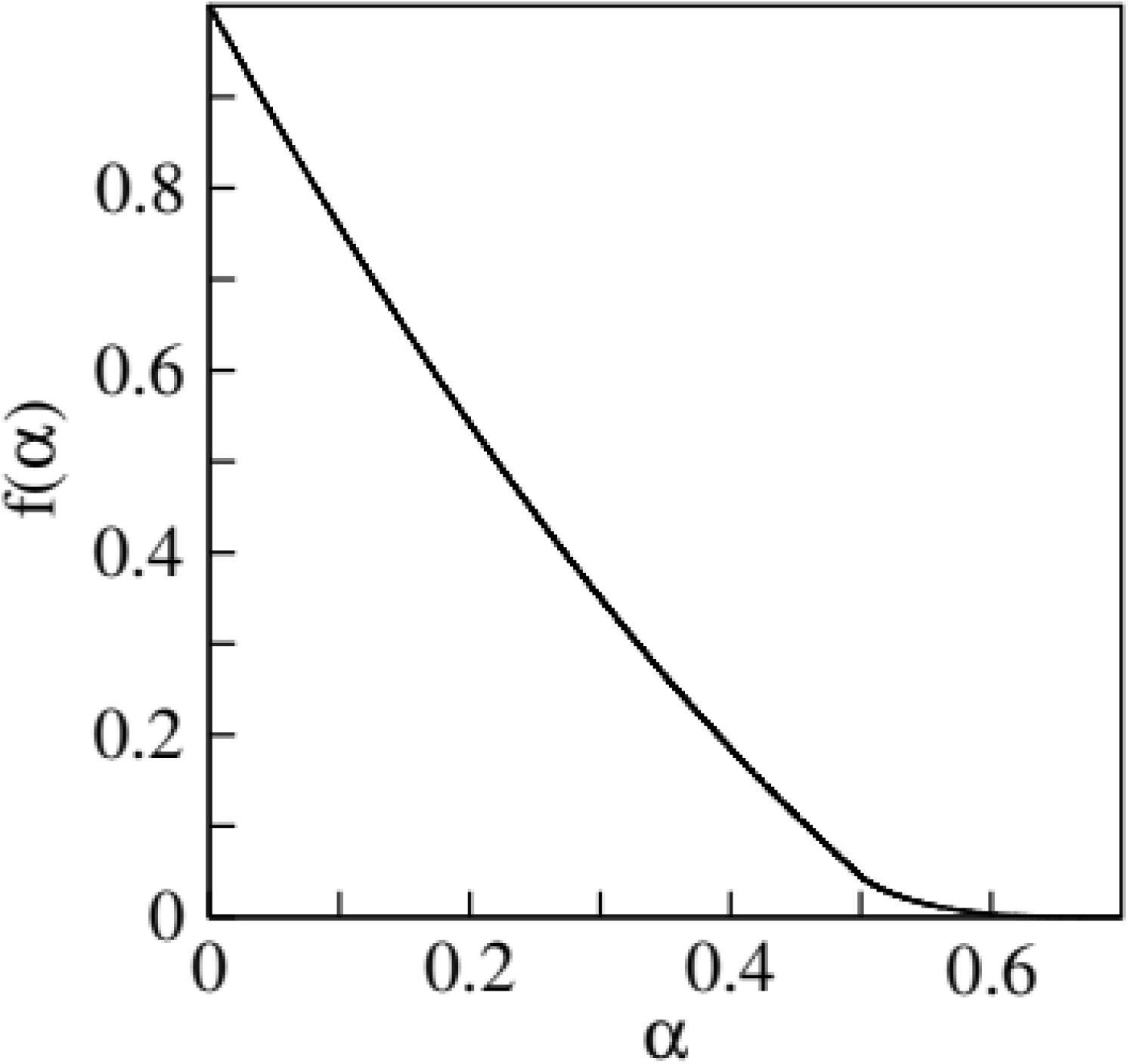
Fraction of the Thon ring modulation, f, at a given spatial frequency recorded on a finite size detector as a function of the ratio of the sideband displacement to the detector size, α, as given in Eq. S6.

## Appendix 3: Model calculation showing Thon Ring fading with defocus for a finite size detector

To model the effects of defocus dependent delocalization on a finite size detector, a 16384×16384 complex array was used. Values in the array outside a circle with diameter of 8192 at the center of the array were set to 1 while the values inside were filled using

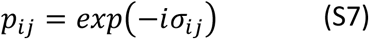

in which σ_ij_ is a random number taken from a Gaussian distribution with standard deviation σ. The effects of defocus and spherical aberration were included by taking the Fourier transform of the whole array and applying the CTF with a chromatic aberration damping envelope. The array was then transformed back to real space, and the center 4096×4096 values extracted. The extracted array was treated in the same way as the experimental images in the main text. Since the extracted array was taken from a larger area, the diffracted side bands are not aliased back into the original image, and the loss of power from sidebands scattered out of the area is compensated by sidebands scattered into the area from outside it.

Since the pixel values are uncorrelated, and the amplitude of the Gaussian distribution is small, the contribution to the power spectra, assuming both sidebands are present, is

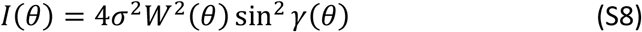

where *γ*(*θ*) is given in Eq.2 and *W*(*θ*) is the amplitude damping envelope. For a detector of finite size, *A*, Eq. S8 becomes

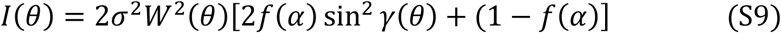

in which *α* = |Δ*ZΘ* − *C_s_θ*^3^|/*A* is the modulus of the ratio of the sideband displacement to the detector size as in defined in Appendix 2, and *f*(*α*) is given in Eq. S6. With a finite size detector, the Thon ring modulation is reduced by *f*(*α*) and sits on an increased background resulting from sideband contributions scattered into the image.

The calculated circularly averaged power spectra for 300 keV electrons at defocus values of 0.5, 1.0, 5.0 and 10.0 μm, as a function of spatial frequency, are shown in Fig S4. In generating these images: σ was set to 0.1, the pixel sampling is 0.5 Å, C_s_ = C_c_ = 2.7 mm, and a full width half maximum energy spread of 0.75 eV was used to generate the damping envelope. Since the Nyquist frequency is at 1 Å^−1^, the value of *θ* corresponding to a given fraction of the Nyquist frequency, *x*, is simply given by *θ* = *λx*.

The calculated sideband displacements as a function of spatial frequency at the different defocus values are shown as red dotted lines in Fig S4. Note that with 0.5 μm defocus as shown in Fig S4a, there is no sideband displacement at *x* = 0.7 corresponding to the Scherzer passband with Δ*Z* = *C_s_θ*^2^.

The Thon ring modulation seen on a finite size detector is sandwiched between lower and upper bounds given by

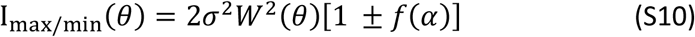

where, as above, α is the ratio of the sideband displacement to the detector size and f(α) is given by Eq. S6. Note that for 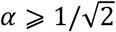 the Thon ring modulation will be totally suppressed *as f*(*α*) = 0. This effect can be clearly seen in Fig. S4(d) where Thon ring modulation is zero for *x* ⩾ 0.79 corresponding to sideband displacements greater than 1448 Å.

The attenuation of the Thon ring modulation with a finite size detector can also be demonstrated by dividing the image from a larger detector into smaller pieces and summing the power spectra from the smaller pieces. This is illustrated in Fig. S5 which shows a simulation similar to those of Fig. S4 but with 2μm defocus. In Fig. S5, the power spectra from the full 4096×4096 image (black) is compared with the sum of the 64 power spectra (blue) obtained from partitioning the image into 512×512 blocks. Both the original and 512×512 blocks were padded to 8192×8192 before taking the FFT. Note that by calculating the power spectra from smaller and smaller areas it is possible to extrapolate to the situation where Thon ring modulation is totally suppressed. By subtracting the resulting power spectra from the original it is therefore possible to extract solely the Thon ring modulation contribution to the power spectra.

**Figure S4.**
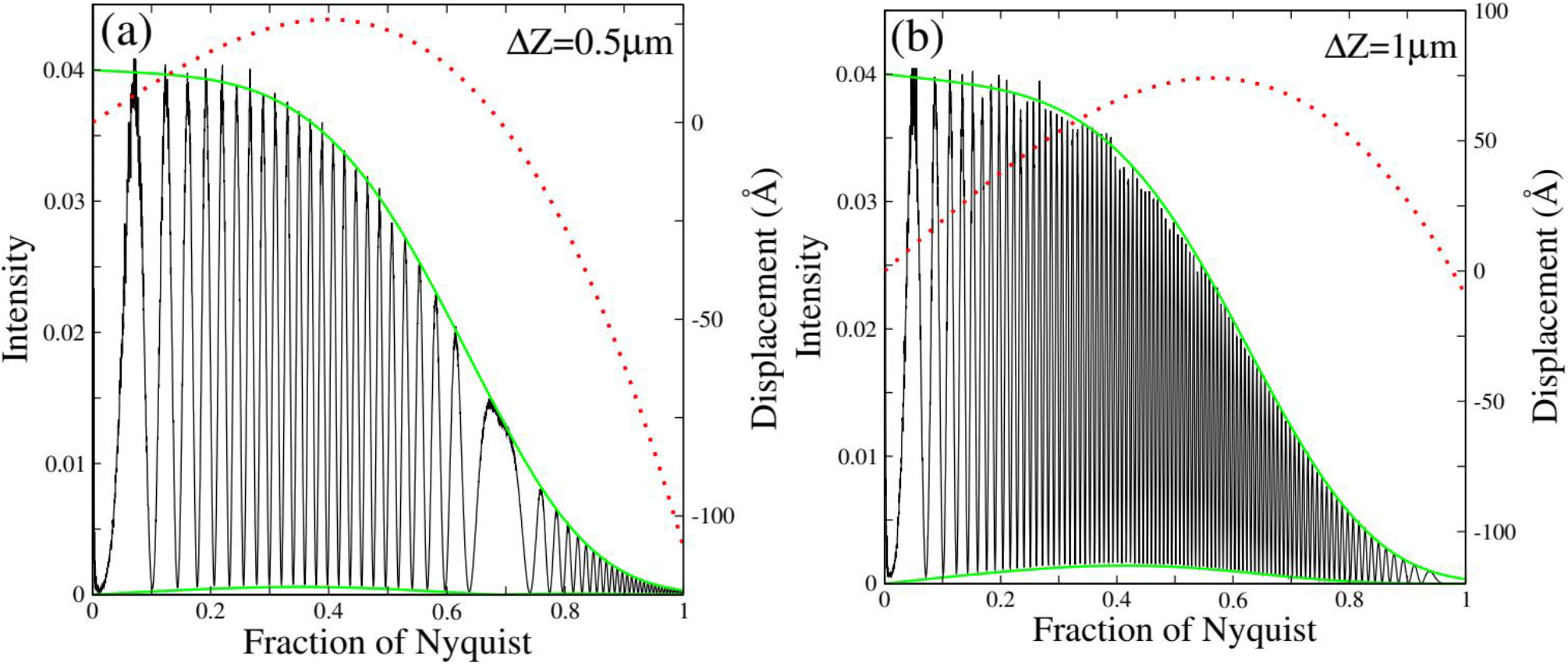
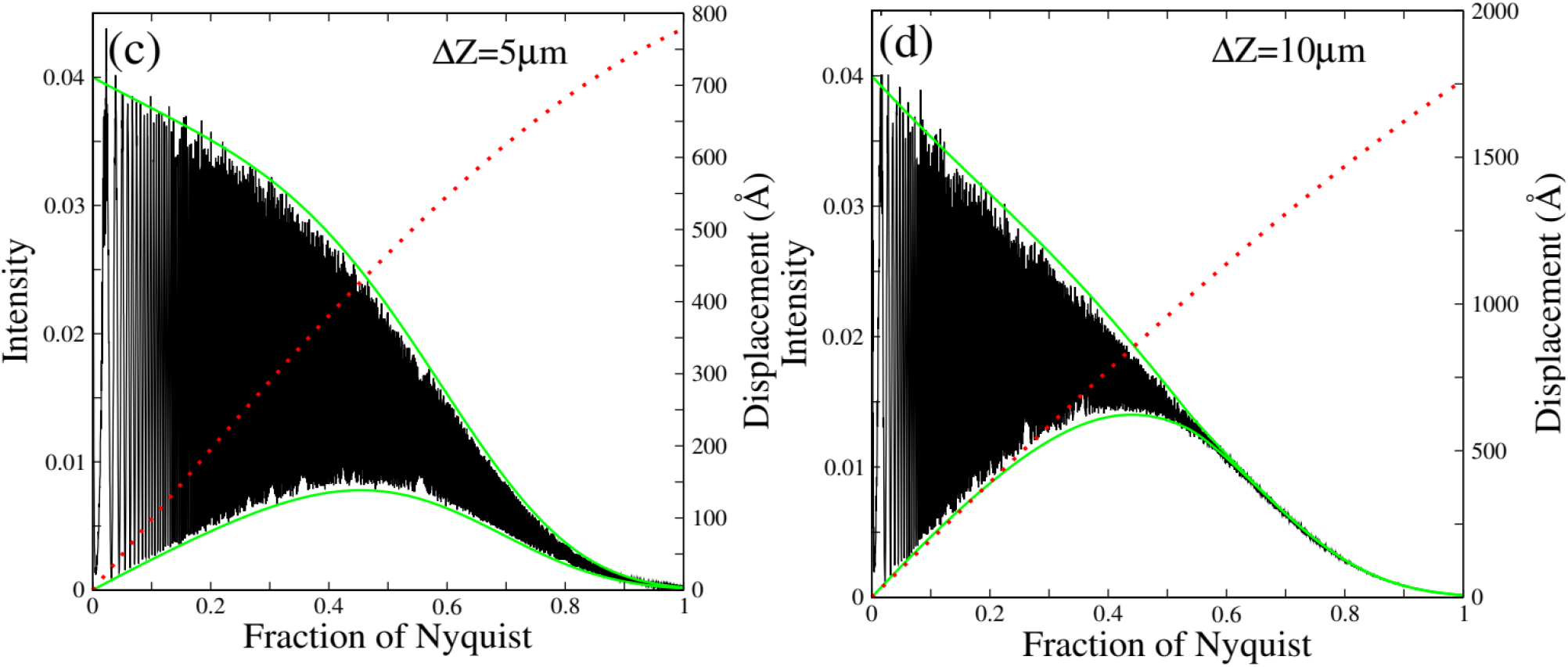
Simulation results for circularly averaged power spectra at different defocus values for 300 keV electrons scattering from a random Gaussian potential as described in the text. Defocus values of 0.5, 1, 5 and 10 μm were used in panels (a) to (d) respectively. In the simulations, a pixel spacing of 0.5 Å was used (giving a Nyquist frequency of 1 Å^−1^), both Cs and Cc were set to 2.7 mm and a temporal damping envelope corresponding to a full width half maximum energy spread of 0.75 eV was used. The upper and lower bounds for the Thon ring modulation calculated using Eq. S10 are shown in green. For each defocus the corresponding sideband displacement as a function of spatial frequency calculated using Eq. S4 is shown (red dotted line) and measured on the right-hand axis.

**Figure S5.**
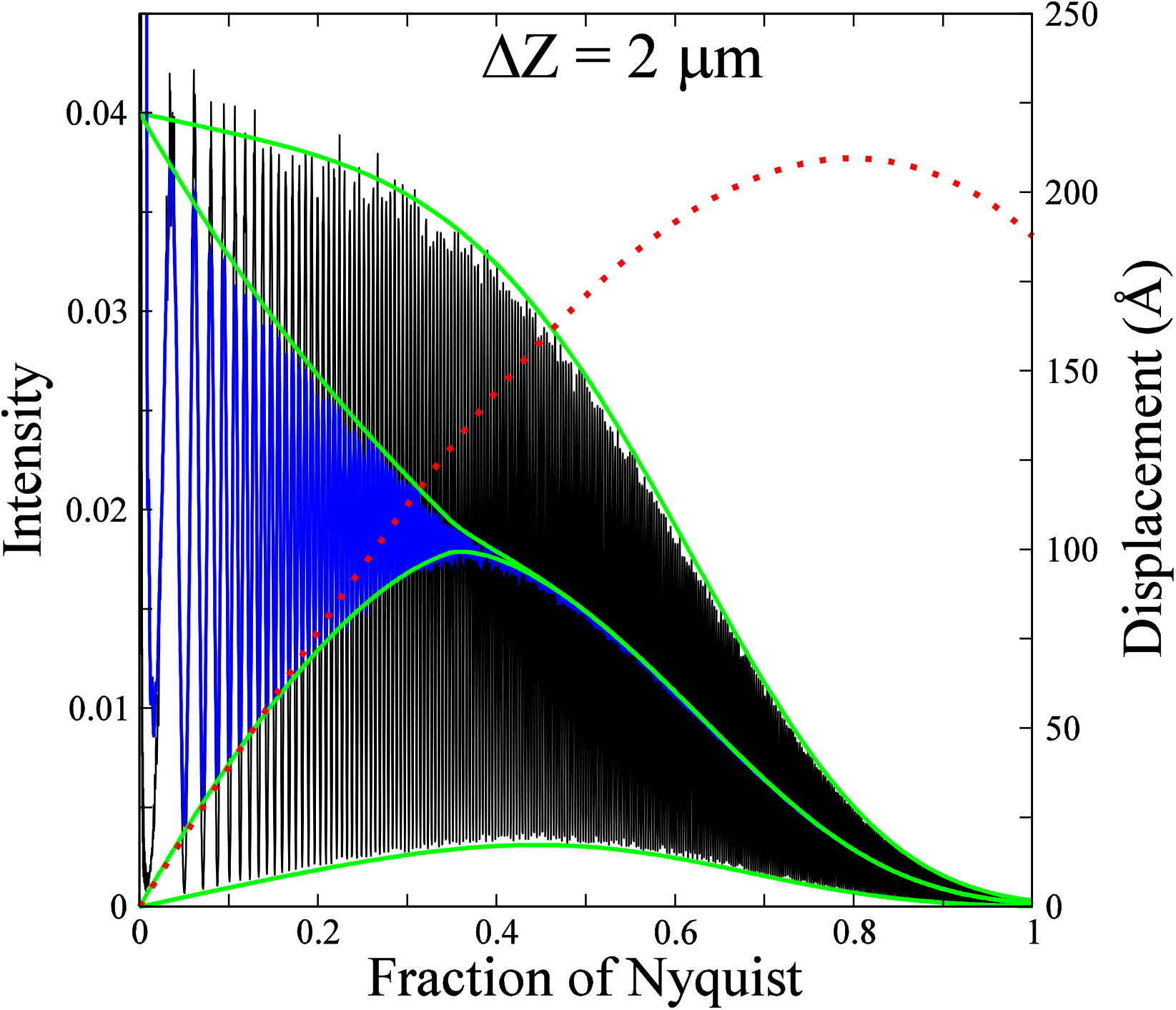
Circularly averaged power spectra as shown in Fig. 4 but for 2 μm defocus. The black curve is calculated using the full 4096 x 4096 area while the blue curve is the sum of the 64 power spectra obtained by dividing the detector into 512×512 blocks. The green lines are the corresponding upper and lower bounds given by Eq. S10. The sideband displacement, a, calculated using Eq. S4, is shown by the dotted red curve and measured on the righthand axis. As a pixel sampling of 0.5 Å has been used, *α* = *a*/256 for the 512×512 blocks. Thus for *a* ⩾ 181 Å, where 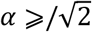 no Thon Ring modulation is expected.

## Appendix 4: Illustrative sideband displacements as a function of electron energy, defocus and spatial resolution

For reference this appendix gives tables for 300 and 100 keV electrons of the sideband displacement at selected defocus and spatial-resolution values calculated using Eq. S4. The longer electron wavelength of 100 keV is reflected in the greater displacements for a given defocus and spatial frequency.

**Table S1:** Sideband displacement as a function of defocus and spatial resolution with 300 keV electrons and Cs of 2.7 mm.

300 keV
| Defocus ($\mu\text{m}$) | Spatial resolution ($\text{\AA}$) | | | | | | |
| --- | --- | --- | --- | --- | --- | --- | --- |
|  | 20 | 10 | 5 | 4 | 3 | 2 | 1 |
| 0.0 | 0.0 | -0.2 | -1.6 | -3.2 | -7.6 | -25.8 | -206.0 |
| 0.5 | 4.9 | 9.6 | 18.0 | 21.4 | 25.2 | 23.5 | -107.6 |
| 1.0 | 9.8 | 19.5 | 37.3 | 46.0 | 58.0 | 72.7 | -9.2 |
| 2.0 | 19.7 | 39.2 | 77.1 | 95.2 | 123.6 | 171.1 | 187.7 |
| 3.0 | 29.5 | 58.9 | 116.5 | 144.4 | 189.2 | 269.6 | 384.6 |
| 4.0 | 39.3 | 78.5 | 155.9 | 193.7 | 254.9 | 368.0 | 581.5 |
| 5.0 | 49.2 | 98.2 | 195.2 | 242.9 | 320.5 | 466.4 | 778.3 |
| 10.0 | 98.4 | 196.7 | 392.1 | 480.0 | 648.6 | 958.6 | 1762.7 |

**Table S2:**
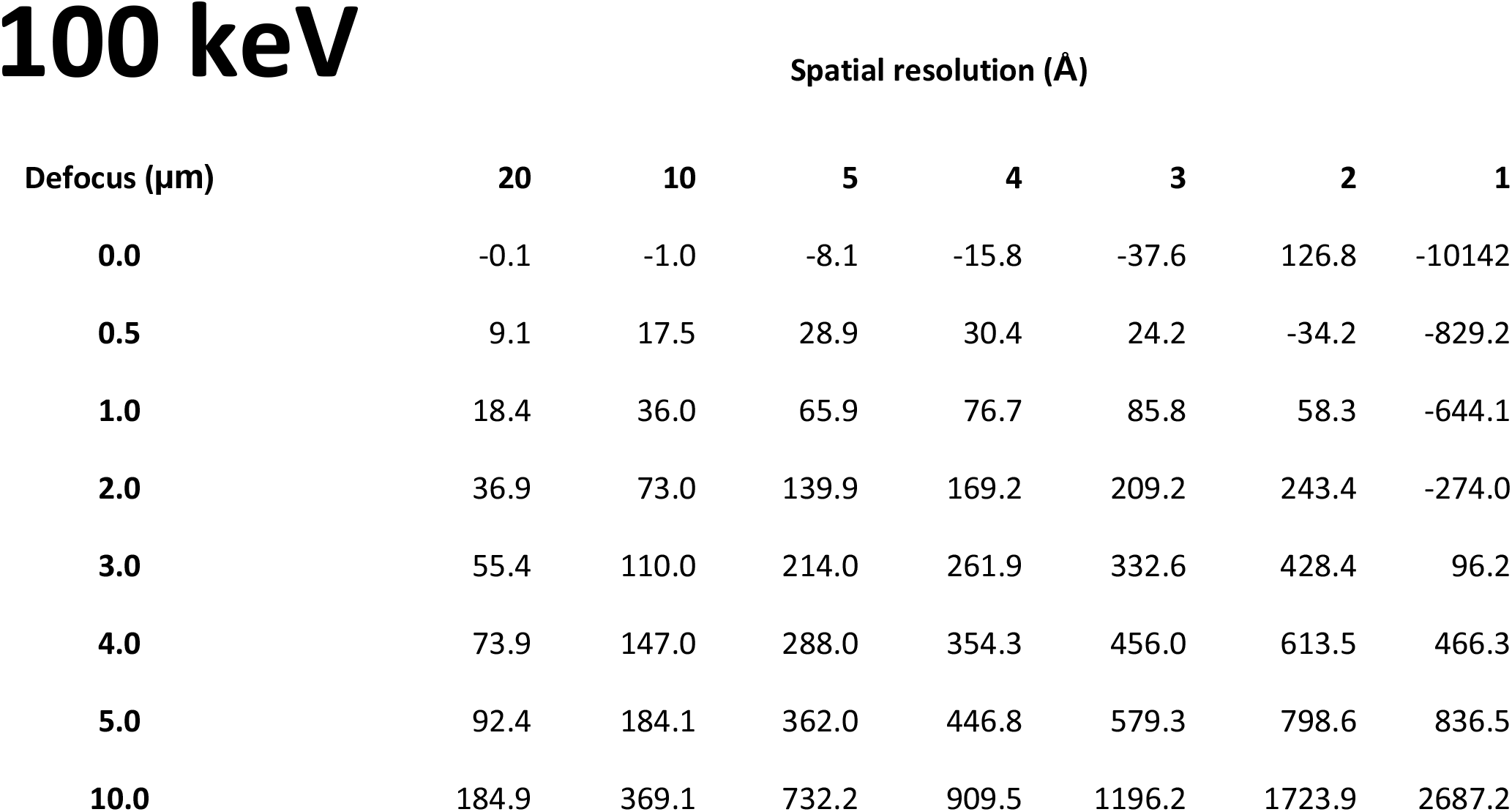
Sideband displacement as a function of defocus and spatial resolution with 100 keV electrons and Cs of 2.0 mm.

